# PhenoGeneRanker: A Tool for Gene Prioritization Using Complete Multiplex Heterogeneous Networks

**DOI:** 10.1101/651000

**Authors:** Cagatay Dursun, Naoki Shimoyama, Mary Shimoyama, Michael Schläppi, Serdar Bozdag

**Author notes:** Permission to make digital or hard copies of part or all of this work for personal or classroom use is granted without fee provided that copies are not made or distributed for profit or commercial advantage and that copies bear this notice and the full citation on the first page. Copyrights for third-party components of this work must be honored. For all other uses, contact the owner/author(s).

## Abstract

Uncovering genotype-phenotype relationships is a fundamental challenge in genomics. Gene prioritization is an important step for this endeavor to make a short manageable list from a list of thousands of genes coming from high-throughput studies. Network propagation methods are promising and state of the art methods for gene prioritization based on the premise that functionally-related genes tend to be close to each other in the biological networks.

In this study, we present PhenoGeneRanker, an improved version of a recently developed network propagation method called Random Walk with Restart on Multiplex Heterogeneous Networks (RWR-MH). PhenoGeneRanker allows multi-layer gene and disease networks. It also calculates empirical p-values of gene ranking using random stratified sampling of genes based on their connectivity degree in the network.

We ran PhenoGeneRanker using multi-omics datasets of rice to effectively prioritize the cold tolerance-related genes. We observed that top genes selected by PhenoGeneRanker were enriched in cold tolerance-related Gene Ontology (GO) terms whereas bottom ranked genes were enriched in general GO terms only. We also observed that top-ranked genes exhibited significant p-values suggesting that their rankings were independent of their degree in the network.

**CCS CONCEPTS:** • Bioinformatics • Biological networks • System biology • Computational genomics

**Availability and implementation:** The source code is available on GitHub at https://github.com/bozdaglab/PhenoGeneRanker under Creative Commons Attribution 4.0 license

## 1. Introduction

Identifying the causal relationship between a gene and complex trait is a challenging problem in functional genomics as their relationship relies on complex and nonlinear interactions of molecular entities [36]. The phenotypic effect of the genotypic aberrations is the result of biological activities that involve the coordinated expression and interaction of proteins or nucleic acids [3]. There are multiple layers of biological processes between genotypic effects to phenotypic outcomes, such as epigenome, transcriptome, proteome that could alter the genotypic effects in many ways.

To represent the multilayered molecular basis of complex traits, biological networks have been utilized extensively [40]. These networks also facilitate data integration, which is useful technique to capture the nonlinear interactions of molecular variations from different layers biological processes while avoiding the limitations and biases of single data types [2, 24]. Each interactome data represent a different aspect of the genotype-phenotype relations. For instance, physical interactome data such as protein-protein interactions (PPI) might have many non-functional PPIs, therefore they are usually complemented by functional interactions [5]. Integrative network models can incorporate datasets from multiple modalities to provide a more comprehensive framework to capture the underlying biology. Analysis of such networks is a powerful approach to demystify the complexity of multilayered molecular interactions and elucidate the genotype-phenotype relations.

Thousands of candidate genes are usually reported to be potentially related to a complex trait by using high-throughput experimental studies such as genome-wide association studies (GWAS). Gene prioritization is essential to shorten a list of thousands of candidate genes into a smaller most probable gene list to facilitate experimental testing [13]. Network propagation methods are promising and state of the art methods for gene prioritization based on the premise that functionally related genes tend to be close to each other in biological networks such as co-expression, PPI and biological pathways [6].

A number of network propagation-based gene prioritization algorithms were previously developed [4, 9, 11, 14, 16, 29, 30, 38]. Among those, random walk with restarts (RWR) algorithms are known to utilize both underlying global network topology and closeness to the known nodes in the network with its restarting property [6]. Although RWR is very effective in gene prioritization, it is known to be biased toward high degree nodes in the network [9].

Recently, a new random walk algorithm called Random Walk with Restarts on Multiplex Heterogeneous Networks (RWR-MH) has been developed [35] as an extension to RWR in heterogeneous networks [18]. RWR-MH performs random walk with restart on a multilayered gene network, which is connected to a single-layer disease similarity network and ranks disease-associated genes based on a set of known disease-associated genes.

Although RWR-MH has multiplex capability for gene networks, it can utilize only one layer of phenotype network. Furthermore, biasedness toward highly connected nodes in the network is a known artifact of the random walk with restart algorithm.

In this study, we developed PhenoGeneRanker, an RWR algorithm on complete multiplex heterogenous networks. PhenoGeneRanker can utilize a heterogeneous network composed of multiplex gene/protein layers as well as multiplex phenotype layers. Also, to account for the biasedness of the RWR algorithm, PhenoGeneRanker generates empirical p-values along with the ranks of genes and phenotypes. To assess the performance of PhenoGeneRanker, we applied it to rank cold tolerance-related genes in rice using a multi-omics rice dataset.

Rice (*Oryza sativa L.*) is the primary source of food for a majority of the world population. Rice is more sensitive to low temperatures than other crops, and cold sensitivity of rice is a major obstacle for its cultivation all around the world [7]. Therefore, understanding the genetic mechanisms of complex traits of rice is important for the world’s food sustainability. It is known that some varieties of rice are more cold tolerant than others [27]. To identify cold tolerance-related genes, a number of Quantitative Trait Loci (QTLs) in rice were identified using GWAS [20, 23, 27, 37, 41]. Several genes were found to be related to the cold response [7].

These genes were involved in signal transduction components, chaperones or transcription factors [31]. To identify larger lists of cold tolerance-related rice genes, we utilized unpublished data from experiments measuring two quantitative phenotypes, Electrolyte Leakage (EL) and Low Temperature Seedling Survivability (LTSS) phenotype experiments using 360 rice cultivars. The results of a companion GWAS study identified thousands of QTL-associated candidate genes [28].

Given the sheer number of candidate cold tolerance-related genes, we applied PhenoGeneRanker to prioritize these genes and rice cultivars. Our workflow is illustrated in Figure 1. We evaluated our results by performing Gene Ontology (GO) enrichment of top and bottom ranked statistically significant candidate genes based on their empirical p-values. Our results showed that the top ranked genes were enriched in cold tolerance-related GO terms, on the other hand the bottom ranked candidate genes were not enriched in cold tolerance-related GO terms. We also observed that in general, top ranked genes had significant p-values suggesting that their rankings were largely based on their closeness to known genes and cultivars rather than solely on their degree in the network. We evaluated our cultivar ranking results based on their cold tolerance classification. We observed that using multiple layers of cultivars as well as an aggregated cultivar layer generated equally-well cultivar ranking based on their cold tolerance classification.

**Figure 1.**
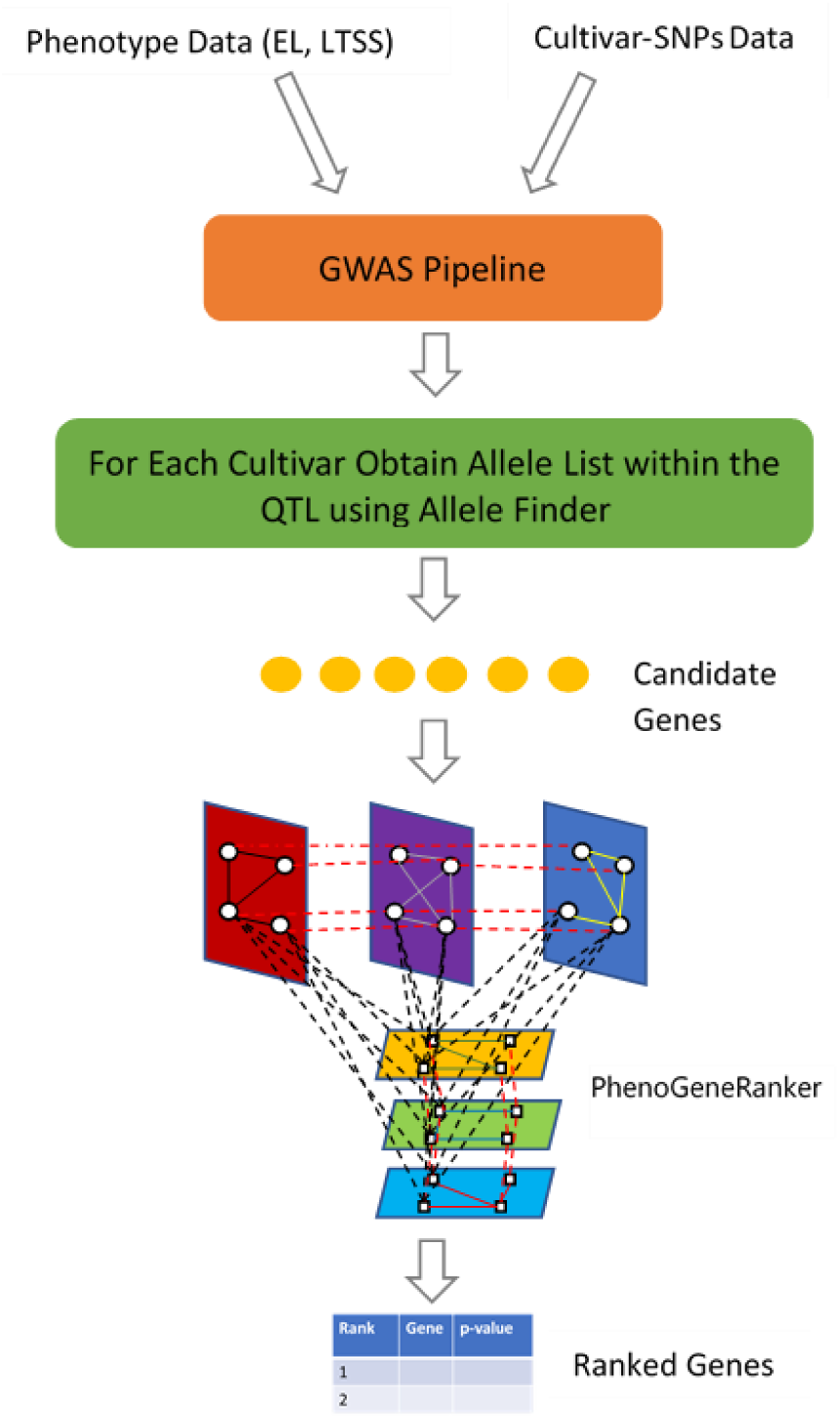
The workflow of identifying and prioritizing cold tolerance-related genes in rice. The first part of the workflow, the GWAS pipeline, to identify candidate genes is described in detail in [28]. Allele Finder was used to obtain allele list [22]. The second phase of gene prioritization was performed by PhenoGeneRanker, the main contribution in the current study. EL: electrolyte leakage; LTSS: low temperature seedling survivability; SNP: single nucleotide polymorphisms; GWAS: genome-wide association study; QTL: quantitative trait loci.

PhenoGeneRanker was implemented in R, and source code can be accessed on GitHub at https://github.com/bozdaglab/PhenoGeneRanker under Creative Commons Attribution 4.0 license.

## 2. Methods

### 2.1. Random Walk with Restart on Complete Multiplex Heterogenous Network (PhenoGeneRanker)

Random walk with restart is a type of network propagation algorithm where the information from pre-specified seed nodes disseminates through the edges of the nodes on the underlying network. Random walk with restart on a heterogenous network was developed to enable random walk by connecting two types of networks namely disease and protein network by establishing bipartite relations between diseases and proteins using phenotype data [18]. RWR-MH was developed to extend this approach by combining multiple protein layers into a multiplex protein network and utilizes the heterogeneous network consisting of bipartite layer and a single-layer disease network [35].

There are several hyperparameters denoted as *r, δ, τ, λ*, and *η* in RWR-MH. *r* is the restart parameter, which controls the probability of jumping to seed nodes during random walk. *δ* is the inter-layer jump probability to controls the probability of staying on the same layer or jumping to another layer. *τ* controls the restart probability to different layers within the multiplex gene network. *η* controls the probability of jumping between gene and phenotype multiplex networks. It has been shown that these hyperparameters have little impact on the rank results [35].

RWR-MH does not have the capability to utilize a multiplex phenotype network. Furthermore, no p-value calculation is perfomed for the ranking results. To address these limiations, in this study, we developed PhenoGeneRanker, which utilizes a complete multiplex heterogeneous network to accommodate two multiplex networks, namely gene/protein and disease/phenotype. In PhenoGeneRanker, complete multiplex heterogeneous network is encoded by a matrix A,

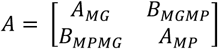

where *A*_*MG*_ is the adjacency matrix of the multiplex gene network, *A*_*MP*_ is the adjacency matrix of the multiplex phenotype network, *B*_*MPMG*_ is the adjacency matrix of the multiplex phenotype-multiplex gene bipartite network, and *B*_*MGMP*_ is the adjacency matrix of the multiplex gene-multiplex phenotype bipartite network (i.e., the transpose of *B*_*MPMG*_).

In addition to all the hyperparameters in RWR-MH, PhenoGeneRanker introduced a new hyperparameter *φ*, which controls restart probability to different layers within the phenotype multiplex network.

#### 2.1.1. Empirical p-value Calculation

Network propagation-based gene prioritization methods are known to be biased toward the high degree nodes in the network [9]. The rank of a node is determined by two criteria: topology of the underlying network and closeness to the seed nodes used for the information propagation. To assess the degree biasedness of each node rank, PhenoGeneRanker employs an empirical p-value calculation based on random seeds. A low p-value suggests that the rank of the node is due to its closeness to the seed nodes and its degree together, whereas a high p-value suggests that the rank of the node is due to its degree rather than its closeness to the seed nodes.

PhenoGeneRanker randomly samples seed nodes using stratified sampling based on the degree of the gene and the cultivar nodes in the network and performs gene prioritization. The number of random seed nodes is set the same as the number of actual gene phenotype seeds. This process is repeated *N* times where *N*=1000 by default. The p-values were calculated based on Eq. 1.

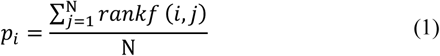

where *rankf*(*i, j*) is an indicator function, and *rankf*(i, j) = 1 if rank of gene *i* for *j*_*th*_ iteration *rank*_*i,j*_ ≤ (*rank*_*i,actual*_ + o*ffset*), and 0 otherwise. *rank*_*i,actual*_ is the rank of gene *i* using actual seeds and we set *offset* to 100 for our calculations. Adding an offset value to the comparison ensures realistic p-values for the top ranked nodes, otherwise it would be biased to get extremely low p-values for the top ranked nodes.

### 2.2. GWAS to Identify Potential Cold Tolerance-Related Genes

To identify genes potentially related to cold tolerance in rice, we utilized the unpublished results of two quantitative phenotypes, percent Electrolyte Leakage (EL) and Low Temperature Seedling Survivability (LTSS) [28, 26]. Cold tolerant rice cultivars have low EL and high LTSS while cold sensitive cultivars have high EL and low LTSS. The results of GWAS analysis based on these phenotypes identified DNA regions that harbor thousands of genes that are potentially related to cold tolerance.

### 2.3. Complete Multiplex Heterogeneous Network for Rice

To prioritize cold tolerance-related genes in rice, we applied PhenoGeneRanker on a complete multiplex heterogenous rice network (Figure 2). We created a three-layer cultivar similarity network namely, EL similarity, LTSS similarity and genotype similarity layers. We also created a three-layer gene interaction network, namely co-expression, PPI and pathway layers. We connected the cultivar multiplex network to the gene network based on the GWAS results. All the layers were composed of undirected and weighted edges. We run PhenoGeneRanker on this complete multiplex heterogenous network using two known cold tolerance-related genes and two cold tolerant cultivars as seeds and ranked all the genes and cultivars. In what follows, we describe each network layer.

**Figure 2.**
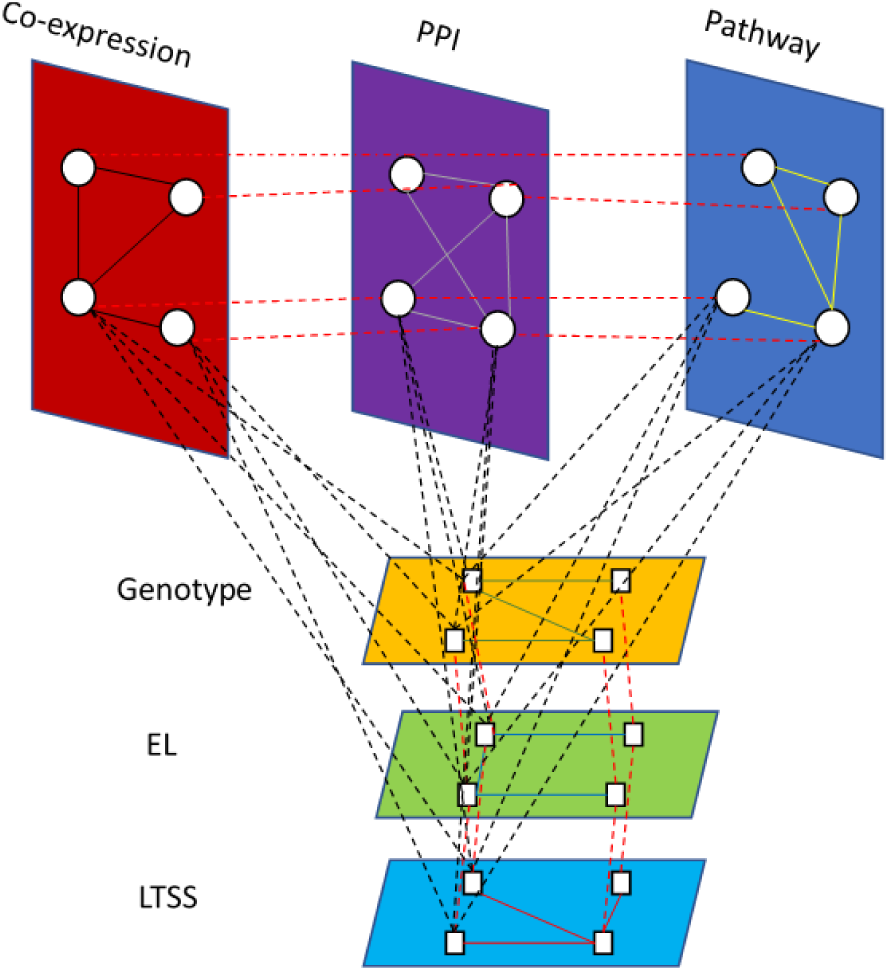
The complete multiplex network structure of PhenoGeneRanker to prioritize cold tolerance-related genes in rice. Layers in the top part form the multiplex gene network. Layers on the bottom forms the multiplex cultivar network. Genes that have relations with phenotypes are connected to phenotypes by black dashed lines. PPI: protein-protein interaction; EL: electrolyte leakage; LTSS: low temperature seedling survivability.

#### 2.3.1. Gene Network

We created a multiplex gene network using co-expression, PPI and pathway layers.

##### 2.3.1.1. PPI Layer

To create the PPI layer, whole physical interactome data for *O. sativa* were downloaded from the STRING V11 database [33]. For protein pairs having multiple interactions between them, we merged their interactions by taking the arithmetic mean of the interaction weights. Protein IDs were mapped to gene IDs by first using alias file in the STRING database and remaining unmapped proteins were mapped using the RAP-DB [25] mapping table. Proteins that were mapped to the same genes were merged into a single node, and the mean of their interaction weights were used as the merged interaction weight.

##### 2.3.1.2. Pathway Layer

To create the pathway layer, *O. sativa* pathway data were downloaded from the Reactome pathway database [10]. We assumed that proteins that co-occur in the same pathways are more similar in function than proteins that occur in different pathways. Given two proteins *i* and *j*, their pathway-based similarity weight was calculated using Eq. 2.

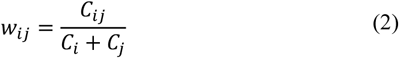

where *w*_*ij*_ is the similarity weight of two proteins *i* and *j, C*_*i*_ and *C*_*j*_ are total number of pathway occurrences of protein *i* and protein *j*, respectively, and *C*_*ij*_is the total number of occurrences of protein *i* and *j* in the same pathway. The protein names were mapped to gene IDs using the R package *rentrez* [39] through NCBI Entrez Gene ID and the RAP-DB mapping table.

##### 2.3.1.3. Co-expression Layer

The co-expression layer was based on the gene expression dataset GSE 57895 in the Gene Expression Omnibus (GEO) database [1]. GSE 57895 has gene expression profiles of shoots and roots in TNG67 (*japonica*) and Taichung Native 1 (*indica*) rice seedlings during cold stress and recovery stages. Specifically, the dataset has expression values for *Control, Cold Treatment 3 hours, Cold Treatment 24 hours*, and *recovery 24 hours* conditions. We filtered the data set to use only Taichung Native 1 shoots, which was one of the cultivars used in the GWAS. We used the R package *WGCNA* to create the co-expression network [17]. Expression value of multiple probes, which were mapped to the same gene was set to the highest value in the group. Absolute values of Pearson’s correlations of the gene expression values were used to create the co-expression network of genes. A soft threshold [15] of power 4 was applied to the co-expression network. To increase the computational efficiency and decrease the number of uninformative correlations, a maximum possible hard threshold value of 0.5724 was applied to shrink the number of edges without losing any nodes in the layer.

To capture the differential impact of cold stress on gene expression values, as an alternative to co-expression layer, we created a differential co-expression layer using *O. sativa* co-expression network from STRING v11 as the reference network. After scaling the co-expression layer obtained using GSE 57895 dataset and the STRING co-expression network to the same interval, we calculated log fold ratio of the intersection of the edges as differential co-expression edge weights.

#### 2.3.2. Cultivar Network

We created a multiplex cultivar network composed of EL, LTSS and genotype layers.

##### 2.3.2.1. EL Layer

The EL layer represents the cultivar similarity network based on EL measurements of cultivars at different temperatures. Dissimilarity values of cultivars were computed by calculating the Euclidian distance of cultivars using average EL values measured at five temperatures. Since EL measurements had high values at some temperatures and might have an effect on similarity, we scaled them to the same interval within each temperature. The dissimilarity values were then converted to similarity values by taking their multiplicative inverse.

##### 2.3.2.2. LTSS Layer

The LTSS layer represents the cultivar similarity network based on the LTSS percentages of cultivars at different temperatures. Dissimilarity values of cultivars were created by calculating the Euclidian distance of cultivars using the average survivability rates measured for five temperatures. As for the EL layer, LTSS rates were scaled to the same interval within each temperature. The dissimilarity values were then converted to similarity values by taking their multiplicative inverse.

##### 2.3.2.3. Genotype Layer

The genotype layer represents the cultivar similarity network based on their common alleles they share. All alleles associated with the 360 cultivars within the QTL regions identified by GWAS were downloaded from Allele Finder [22]. We used the alleles (homozygous and heterozygous) from the coding regions only based on the assumption that those alleles would have more impact on the phenotype than non-coding alleles. Dissimilarity between each cultivar pair was calculated using the Gower distance method [12] implemented in the R package *cluster* [21]. Dissimilarity values were then converted into similarity values as 1 − Gower Distance.

#### 2.3.3. Gene-Cultivar Bipartite Layer

The gene-cultivar bipartite layer connects the cultivar nodes to the candidate gene nodes (i.e., the genes reported in the GWAS study). The similarity score (i.e., the weight of the edge) between a cultivar and a gene was calculated based on the number of homozygous alleles in the coding region of this gene for this cultivar. We considered only the homozygous alleles because we assumed they would have stronger phenotypic effects than the heterozygous alleles. To compute the similarity scores, for each gene, the homozygous allele counts for each cultivar were log transformed and normalized by the total number of homozygous alleles across all the cultivars. The resultant similarity values were used as the gene-cultivar bipartite layer.

### 2.4. Seeds

Seeds are information sources for the network propagation algorithms. The RWR algorithm returns to the seed nodes at each restart. We used two genes as seeds for our approach; LOC_Os06g39750 and LOC_Os09g29820. LOC_Os06g39750 was reported as a candidate gene for the qPSST6 QTL, which controls the percent seed set under cold water treatment [32]. LOC_Os09g29820 encodes the transcription factor bZIP73 and has allelic differences between cold tolerant *japonica* and cold sensitive *indica* cultivars. It was reported to differentially regulate the expression of cold tolerance-related genes for rice because it has facilitated rice subspecies *japonica* cold climate adaptation [19].

To determine the cultivar seeds, we sorted the cultivars based on lowest EL and highest LTSS and LT50 (i.e. the temperature at which 50% of seedlings died) values and selected the top two cultivars as the cultivar seeds.

## 3. Results

In this study, we developed a gene prioritization tool, PhenoGeneRanker that uses multiplex gene and phenotype networks to prioritize disease-associated genes. To assess PhenoGeneRanker, we applied it on a rice dataset to rank cold tolerance-related genes. In this system, cold sensitivity is the disease and cold tolerance is the non-disease state. We used candidate cold tolerance-related genes identified by EL and LTSS GWAS experiments using 360 rice cultivars [28]. We obtained 10,880 candidate cold tolerance-related genes within QTL regions identified by GWAS QTLs [28]. A complete multiplex gene network was generated using co-expression/differential co-expression, PPI and pathway layers. A complete multiplex cultivar network was generated using EL, LTSS and genotype layers. In the following sections, we use the word “complete” for the multiplex networks that use all possible layers.

We built the gene network layers using publicly available expression, pathway and PPI datasets (see Methods). For the gene network, we had two expression-based layers, namely co-expression and differential co-expression. Only one of these layers was used in a given final network. The number of nodes and edges of each layer in the final complete multiplex heterogenous network is shown in Table 1. For the gene network, the pathway layer was the smallest and co-expression was the largest layer. In the cultivar network, each layer was a fully connected network as we were able to compute cultivar similarity for all pairs. Among 10,880 candidate genes, only 3,913 were represented in the gene network as there was no gene level data available for the other genes. When differential co-expression layer was used instead of co-expression layer, 2,924 of the candidate genes were represented in the network.

**Table 1.** The numbers of nodes and edges in each layer of the final complete multiplex rice network.

| Layer | Number of Nodes | Number of Edges |
| --- | --- | --- |
| PPI | 13,007 | 667,580 |
| Pathway | 2,585 | 140,042 |
| Co-expression | 18,402 | 7,023,448 |
| Differential co-expression | 11,624 | 190,278 |
| EL | 360 | 64,620 |
| LTSS | 360 | 64,620 |
| GT | 360 | 64,620 |

We set the parameters for all reported results as follows; r = 0.7, δ = 0.5, *η* = 0.5, λ = 0.5 We set τ and φ parameter values to 0.5 to give equal probability of restarting to each gene and cultivar layers, respectively within the multiplex networks.

We generated multiple gene/cultivar rankings using combinations of the network layers and seeds to analyze their effects on the prioritization of genes/cultivars.

### 3.1. Effects of the Gene Layers and the Cultivar Seeds on the Gene Rankings

We analyzed the effects of the gene layers and the cultivar seeds to evaluate their individual contributions to the gene rankings. We compared the different gene ranking results by using Kendall’s tau coefficient, which compares pairs of ranks by evaluating their concordance of the two ranks. Kendall’s tau coefficient between two ranks are 1 if the ranks are identical and −1 when one rank is the inverse of the other. We observed that each gene layer had a large effect on the gene rankings suggesting that integration of different types of datasets contributed significantly to the results (Table 2). For the ranking of top 200 genes, the largest contribution came from the co-expression layer, followed by PPI and pathway layers.

**Table 2.** Kendall’s tau coefficients of the gene rankings using different gene layers and/or seeds. Gene ranking obtained after modification was compared with the reference gene ranking. The reference gene ranking was generated by running PhenoGeneRanker on the complete multiplex heterogenous network with co-expression layer using the gene and cultivar pair seeds. Whole Rank Tau shows the comparison of the entire gene ranking in the gene network. Top 200 Common shows the number of common ranked genes in top 200. Top 200 Tau shows the comparison of the ranking of the top 200 common genes.

| Network/Seed Modification | Whole Rank Tau | Top 200 Common | Top 200 Tau |
| --- | --- | --- | --- |
| Not using the cultivar seeds | 0.618 | 50 | 0.667 |
| Removing the pathway layer | 0.892 | 182 | 0.905 |
| Removing the PPI Layer | 0.684 | 180 | 0.873 |
| Removing the co-expression layer | 0.486 | 36 | 0.924 |
| Using the differential co-expression layer instead of the co-expression layer | 0.694 | 33 | 0.708 |

The contribution of the cultivar seeds to the rankings was important since we incorporated the cultivar phenotype and genotype similarity information through the use of cultivar seeds. As shown in Table 2 that the usage of the cultivar seeds with the gene seeds changed the gene rankings substantially.

### 3.2. Effects of the Cultivar Layers on the Cultivar Rankings

We analyzed the effects of the cultivar layers to evaluate their individual contribution to the cultivar rankings. We computed Kendall’s tau coefficient between the cultivar rankings obtained using the complete multiplex heterogeneous network with co-expression layer and the cultivar rankings by removing single cultivar layers, using a single aggregated cultivar layer or a random cultivar layer. We created the aggregated cultivar layer by taking geometric means of common edges of all cultivar layers and resultant edge values of the layer were rescaled to the same interval with other layers. The random cultivar layer was generated as fully connected cultivar network like the other cultivar layers where edge values were generated from standard normal distribution and scaled to same interval with other layers. The results showed that the largest effect occurred when the LTSS layer was removed (Table 3). We also observed that the aggregated layer result was the most similar to the result of the complete multiplex heterogeneous network. As expected, the cultivar rankings based on a random cultivar layer was quite different.

**Table 3.**
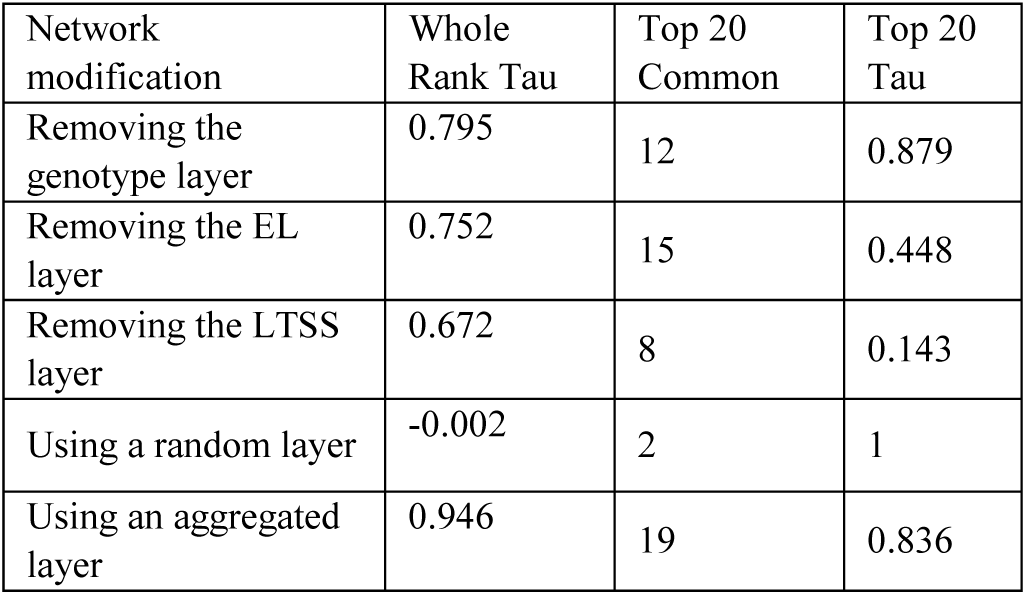
Kendall’s tau coefficients of the cultivar rankings using different cultivar layers. Cultivar ranking obtained after modification was compared with the reference cultivar ranking. The reference cultivar ranking was generated by running PhenoGeneRanker on the complete multiplex heterogenous network with co-expression layer using the gene and cultivar pair seeds. Whole Rank Tau shows the comparison of the entire cultivar ranking in the cultivar network. Top 20 Common shows the number of common ranked cultivar in top 20. Top 20 Tau shows the comparison of the ranking of the top 20 common cultivars.

| Network modification | Whole Rank Tau | Top 20 Common | Top 20 Tau |
| --- | --- | --- | --- |
| Removing the genotype layer | 0.795 | 12 | 0.879 |
| Removing the EL layer | 0.752 | 15 | 0.448 |
| Removing the LTSS layer | 0.672 | 8 | 0.143 |
| Using a random layer | -0.002 | 2 | 1 |
| Using an aggregated layer | 0.946 | 19 | 0.836 |

We further analyzed the cultivar rankings described in Table 3 by overlaying the cultivar cold tolerance classifications, namely *tolerant, intermediate* and *sensitive*. These classifications were based on the clustering of admixed and *aromatic* cultivars as *intermediate* between cold *tolerant* cultivars (*temperate japonica* and *tropical japonica*, i.e. low EL and high LTSS) and cold *sensitive* cultivars (*aus* and *indica*, i.e. high EL and low LTSS) using principal component analyses (PCA; [26]). There were 207 tolerant, 20 intermediate and 131 sensitive cultivars (Figure 3). Since two of the cultivars were seeds, they were not shown in the ranking. As shown clearly in Figure 3, integration of data contributes positively to separation of the cultivars by their classes. As expected, cultivar ranking obtained by only a random cultivar layer resulted in a random ranking of the cultivars based on their classes.

**Figure 3.**
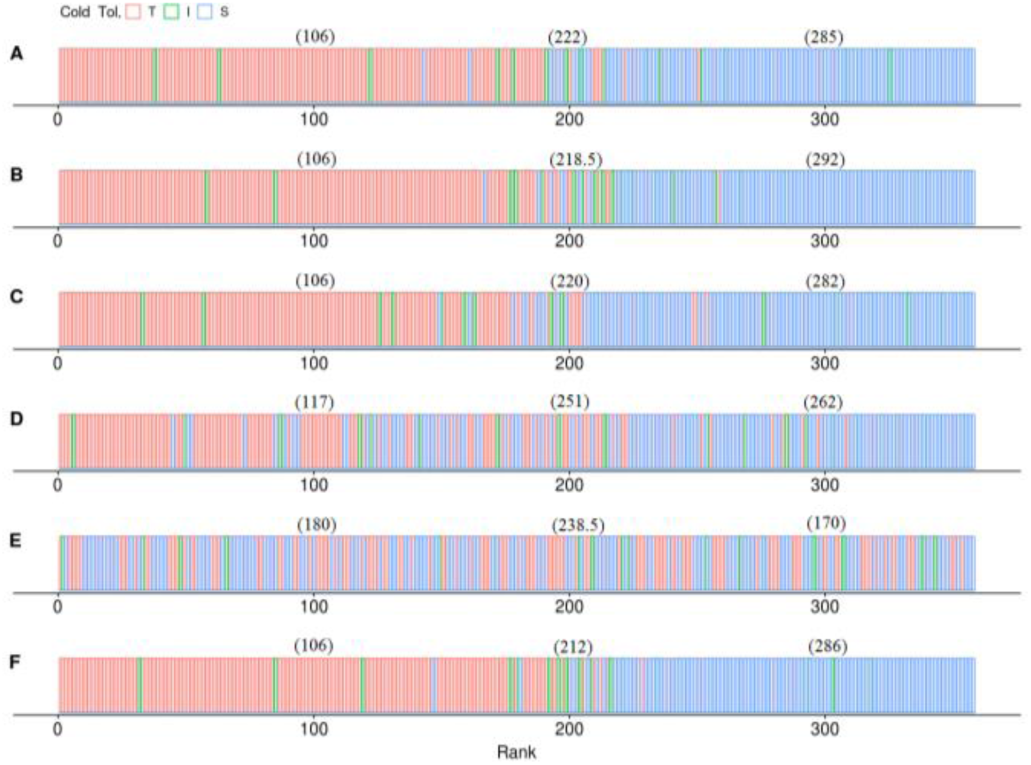
The cultivar ranking results. In each PhenoGeneRanker run, gene and cultivar pair seeds were used on a multiplex network where gene network consisted of co-expression, PPI and pathway layers and cultivar network consisted of A: EL, LTSS and genotype layers. B: EL and LTSS layers only. C: LTSS and genotype layers only. D: EL and genotype layers only. E: A single random cultivar layer. F: Single aggregated layer using EL, LTSS and genotype. The median rank of each cold tolerance classification are shown in parentheses for each result in the order of cold tolerant, intermediate and cold sensitive. T: cold tolerance, I: intermediate, S: sensitive.

In Figure 3, median ranks of cultivars by their cold tolerance classification for different layer combinations are shown in paratheses for each result. In a perfect ranking where each cultivar class is clustered together, the median ranking of tolerant, intermediate and sensitive cultivars would be 104, 217 and 293, respectively. As shown in Figure 3, the complete multiplex layer and the aggregated layer had quite similar results. We particularly observed that cold tolerant and sensitive cultivars were quite similar when using complete multiplex network and aggregated layer. On the other hand, the ranking of intermediate cultivars was slightly different and equally close to the expected median ranking. Interestingly when we removed the genotype layer the median ranks became closer to expected values. Moreover, not using LTSS layer had substantial negative effect on the median ranks. This result suggests that the LTSS is the most informative cultivar layer.

### 3.3. The Top Ranked Rice Genes were Enriched in Cold Tolerance-Related GO Terms

To assess the functional relevance of the top ranked rice genes by PhenoGeneRanker, we used agriGOv2 [34] to perform GO enrichment of the top 200 ranked all and candidate-only genes in multiple results. For these GO enrichments, we only considered the genes whose empirical ranking p-value ≤ 0.05. As a negative control, we also performed GO enrichment for the bottom 200 ranked candidate rice genes. We used Fisher as test method and Bonferroni as multiple test correction method with significance cutoff value of 0.05.

Enrichment results for the top 200 genes, regardless of whether they were candidate or not, are shown in Figure 4, and GO enrichment results for the top 200 candidate genes are shown in Figure 5. Both GO enrichment analysis resulted similar GO terms that were previously identified as part of cold tolerance strategies of rice cultivars [31]. The top 200 candidate gene GO term enrichment analysis suggests highly significant candidate genes are involved in lipid metabolism (GO:0006629, GO:0006631, GO:0006633, GO:0008610, GO:0016020, and GO:0044255), carboxylic acid metabolism (GO:0006082, GO:0016053, GO:0019752, GO:0032787, GO:0042180, GO:0043436, GO:0044281, GO:0044283, and GO:0046394), acyl transferase activity (GO:0016746 and GO:0016747), and oxidation-reduction process (GO:0016491 and GO:0055114). Interestingly, the two most significant clusters of GO terms (lipid metabolism, and carboxylic acid metabolism) have child terms relevant to known cold stress associated plant hormones including abscisic acid, gibberellin, jasmonic acid, and salicylic acid. Previous studies have shown a significant involvement of plant hormone pathways in regulating cold stress response and tolerance [8]. Moreover, lipid metabolism and oxidation-reduction process suggest genetic pathways that might protect plasma membrane integrity via phospholipid biosynthesis and scavenging reactive oxygen species, which would result in low EL and high LTSS.

**Figure 4.**
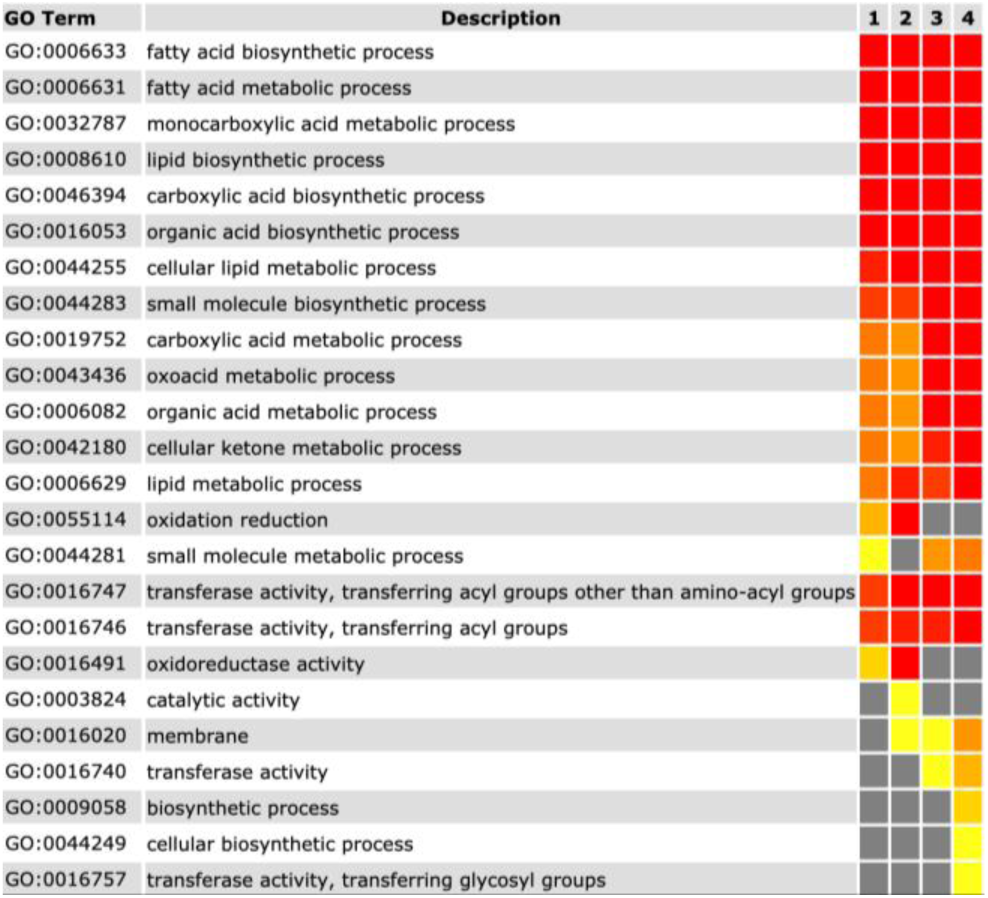
GO enrichment results using statistically significant ranked genes within the top 200 in four different results. All results were generated using the complete multiplex gene and complete multiplex cultivar networks. All genes in the network were used as background. 1: Only gene seeds, co-expression layer. There were 199 statistically significant genes. 2: Only gene seeds, differential co-expression layer. There were 190 statistically significant genes. 3: Gene and cultivar seeds, co-expression layer. There were 80 statistically significant genes. 4: Gene and cultivar seeds, differential co-expression layer. There were 46 statistically significant genes. Only significant GO terms are reported; yellow to red color indicates increase of statistical significance, gray color indicates non-significance (i.e., p-value > 0.05).

**Figure 5.**
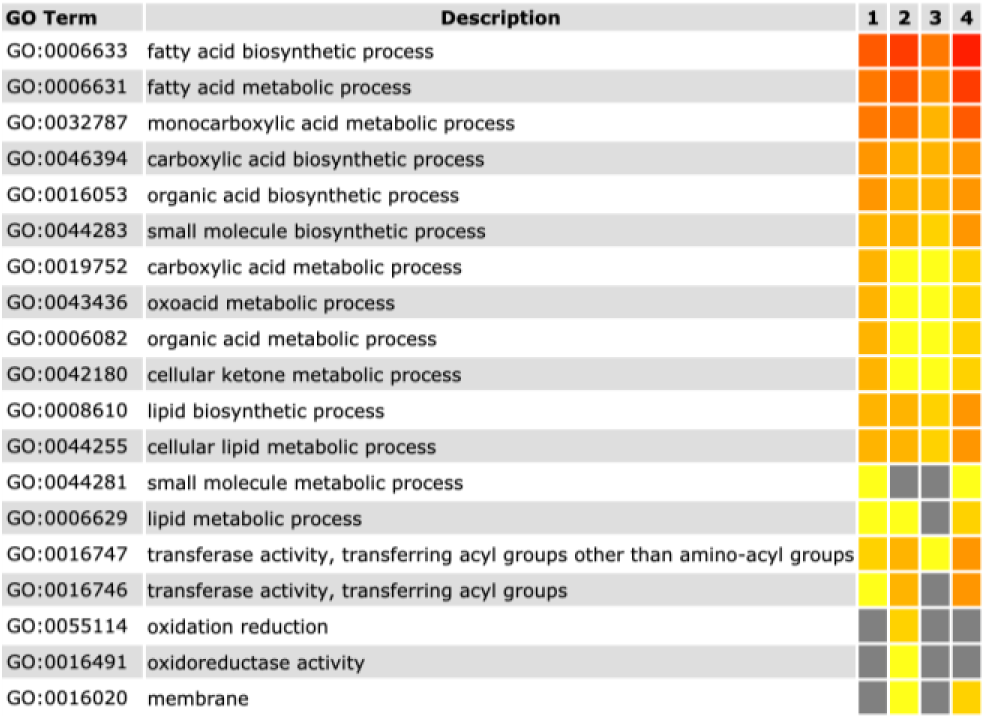
GO enrichment results using statistically significant ranked candidate genes within the top 200 in four different results. All results were generated using the complete multiplex gene and complete multiplex cultivar networks. All candidate genes in the network were used as background. 1: Only gene seeds, co-expression layer. There were 37 statistically significant candidate genes. 2: Only gene seeds, differential co-expression layer. There were 28 statistically significant candidate genes. 3: Gene and cultivar seeds, co-expression layer. There were 66 statistically significant candidate genes. 4: Gene and cultivar seeds, differential co-expression layer. There were 17 statistically significant candidate genes. Only significant GO terms are reported; yellow to red color indicates increase of statistical significance, gray color indicates non-significance (i.e., p-value > 0.05).

By contrast, GO enrichment for the bottom 200 genes (Figure 6) when using gene pair as seeds resulted in identifying these genes as associated with the basic process of DNA replication. Taken together, all terms seem to be tied to different components of the DNA replication including: helicase activity, nucleic acid binding, ribonuclease activity, and RNA binding. While these genetic pathways may still play a role in cold tolerance mechanisms, it is more likely that they are involved in basic biological functions rather than specific cold stress response pathways. The results obtained using gene and cultivar pair as seeds (i.e., result #3 and #4 in Figure 6) had zero and one enriched GO term, respectively. This suggests that using gene and cultivar pairs perform effectively in moving non-cold tolerance-related genes to the bottom of the gene ranking.

**Figure 6.**
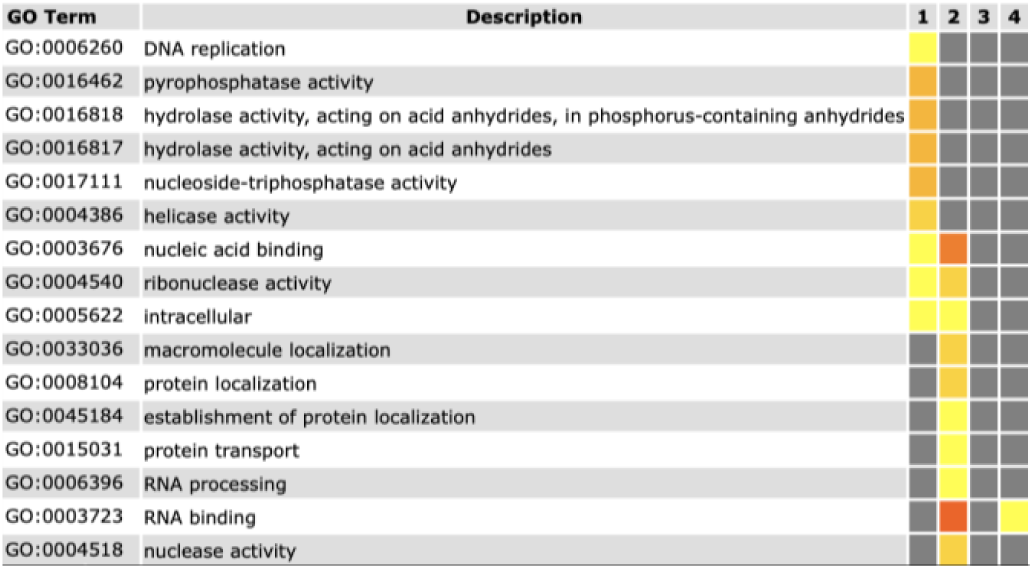
GO enrichment results using bottom 200 candidate genes in four different rank results. All results were generated using complete multiplex gene and multiplex cultivar networks. 1: Only gene seeds, co-expression layer. 2: Only gene seeds, differential co-expression layer. 3: Gene and cultivar seeds, co-expression layer. 4: Gene and cultivar seeds, differential co-expression layer. Only significant GO terms are reported; yellow to red color indicates increase of statistical significance, gray color indicates non-significance (i.e., p-value > 0.05).

### 3.4. Top Ranked Genes Tend to Have Lower p-values

PhenoGeneRanker employs a random stratified sampling to calculate empirical p-values for the rankings (see Section **Error! Reference source not found.**). The relationship between empirical p-value and ranking of candidate genes is shown in Figure 7. Low p-values suggests that the rank of a gene does not solely depend on the topology of the underlying network, but also on its closeness to the cold tolerant gene and cultivar seeds. We observed that p-values assigned to the top-ranked genes were generally low, and p-values for higher ranked genes were higher. We observed that high degree genes tend to be clustered in the top of the ranking. However, certain high degree genes have high p-values suggesting that PhenoGeneRanker is not biased to the high degree genes.

**Figure 7.**
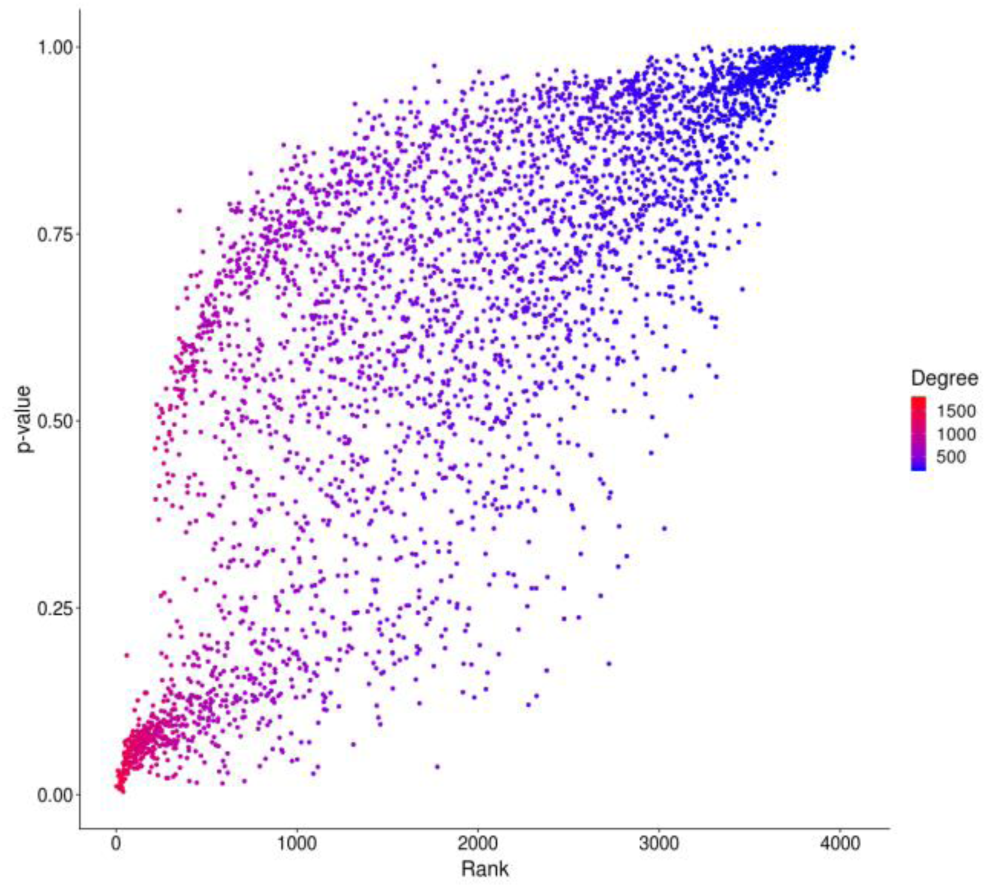
The scatterplot between p-values and gene ranking of the candidate genes. p-values were obtained from the result where complete multiplex heterogenous network with co-expression layer was used with gene and cultivar pair seeds.

## 4. Discussion

In this study, we developed a gene prioritization tool called PhenoGeneRanker that integrates gene and phenotype-based datasets in a multiplex heterogenous network. PhenoGeneRanker extends RWR-RH by enabling multiplex phenotype layers and computing empirical p-values of the ranking.

We applied PhenoGeneRanker on a rice dataset to prioritize cold tolerance-related rice genes. We identified potential cold tolerance-related genes that were ranked lowest with a statistically significant p-value. GO enrichment of the top 200 of the results showed that the lowest ranked genes were enriched in lipid metabolism and fatty acid synthesis GO terms (Figure 4 and Figure 5). Those terms were previously shown to play an important role in cold tolerance of *O. sativa* [31].

We analyzed the effect of the network layers on the results by comparing Kendall’s tau coefficients of the ranks from multiple results. Adding different gene layers had significant impact on the results. Co-expression, PPI and pathway layers had the largest effects in decreasing order (Table 2). Since a large network size implies more information, effects of layers on the results were proportional to sizes of the layers. We observed that the use of cultivar seeds with gene seeds had significant effect on the ranks (Table 2). We also observed that cultivar layers affected the rank of cultivars (Table 3). However, since all cultivar layers were fully connected networks, their individual impacts were not as significant as gene layers. The LTSS layer had the largest impact on the cultivar rankings.

To better recapitulate the transcriptional response signals to cold tolerance, we utilized a differential co-expression layer as an alternative to co-expression layer. However, the reference co-expression network obtained from STRING database had some limitation. The reference co-expression network did not have information about the growth stage and the tissue source of the expression data. However, our GWAS results were based on the early growth stage of rice cultivars, and phenotypic measurements were made on samples using only shoots and leaves. A differential co-expression layer with a more relevant reference co-expression network would potentially be more useful.

To increase the reliability of our results we generated empirical p-values for our PhenoGeneRanker rank results by stratified sampling based on the degree of the nodes in the network. We observed that low ranked genes had overall lower p-values, which shows closeness of the ranked gene to the seeds rather than sole centrality in the underlying network topology (Figure 7).

One of the unique features of PhenoGeneRanker is to allow multiplex layers for the phenotype network. Multiplex networks were shown to have better performance than aggregated monoplex networks [35]. Using this feature, different from previous studies, we were able to integrate phenotype (EL and LTSS) and genotype information in a multiplex cultivar network. We tested this extension by comparing results using multiplex and aggregated cultivar networks and observed that the cultivar rankings obtained by using these networks were similar (Table 3, Figure 3). The multiplex phenotype network feature could be effectively utilized in cases where network size is bigger and have complex structure which is usually the case for studies such as drug-gene interaction studies.

## ACKNOWLEDGMENTS

This study was funded in part by grant HL64541 from the National Heart, Lung, and Blood Institute on behalf of the NIH (to M. Shimoyama) and USDA-AFRI-NIFA grant @2016-6701302487 and #2018-67014027470 (to M. Schläppi).

